# Dnmbp interacts with Daam1 to facilitate assembly of cadherin-mediated junctions in epithelializing nephric tubules

**DOI:** 10.64898/2026.07.02.736208

**Authors:** Brandy L. Walker, Bridget D. De Lay, Yogesh Srivastava, Mark E. Corkins, Vanja Krneta-Stankic, Adrian Romero, Rachel K. Miller

## Abstract

The mature kidney contains approximately one million nephrons, and defects arising during nephron development can result in lifelong renal impairment, often culminating in kidney failure and transplantation. Nephric tubule formation requires coordinated epithelial processes, including polarity, adhesion, signaling, and vesicle transport; however, how these processes are integrated during kidney morphogenesis remains unclear. Dynamin binding protein (Dnmbp) is a multi-domain scaffolding protein expressed in human kidneys that is involved in several cellular processes. Using the *Xenopus* embryonic kidney, we previously demonstrated that Dnmbp is essential for nephrogenesis, yet the mechanisms by which it influences nephron development remain undefined. Here, we identify Dnmbp as a novel interacting partner of the Wnt/planar cell polarity effector Daam1. The interaction between Daam1 and Dnmbp was independently identified in two yeast two-hybrid screens, biochemically verified, and supported by structural modeling predictions of a Daam1-Dnmbp complex. In developing *Xenopus laevis* kidneys, Dnmbp localized to punctate structures associated with E-cadherin-rich cell-cell contacts. Dnmbp depletion significantly reduced junctional E-cadherin localization in both epithelializing and mature nephric tubules without affecting total E-cadherin levels, indicating a role in E-cadherin recruitment or stabilization at adherens junctions. Furthermore, expression of human *DNMBP* rescued the junctional defects, confirming the specificity of the loss-of-function phenotype. Together, these findings identify Dnmbp as an essential regulator of kidney development and support a model in which Dnmbp provides a mechanistic link between Wnt/PCP signaling, Cdc42 activation, and adherens junction formation during nephrogenesis.

## INTRODUCTION

The formation of a functional nephron requires the precise coordination of epithelial polarity, cell-cell adhesion, cytoskeletal dynamics, and vesicular transport. These processes collectively establish spatial organization, mechanical integrity, and directional behaviors that allow nephron progenitor cells to assemble into a cohesive, elongated tubule. (Dressler, 2006; Little and McMahon, 2012; Bryant and Mostov, 2008) During nephron epithelialization, the formation of stable cadherin-mediated adherens junctions is essential for establishing and maintaining cell-cell contacts and cytoskeletal intercellular connections, as well as for facilitating polarized cellular rearrangements that drive nephron morphogenesis (Costantini and Kopan, 2010; Nishimura and Takeichi, 2009). In epithelial tissues, adherens junctions and polarity complexes are also closely integrated with intracellular membrane trafficking pathways that control the delivery and recycling of junctional and polarity proteins (Bryant and Mostov, 2008; Rodriguez-Boulan and Macara, 2014). Although the molecular machinery governing polarity and adhesion has been extensively characterized, the mechanisms that integrate vesicular transport with cytoskeletal remodeling during nephron morphogenesis remain poorly understood (Bryant and Mostov, 2008; Datta et al., 2011).

Current models of renal development indicate that non-canonical Wnt/planar cell polarity (PCP) signaling regulates differentiation and epithelialization (Miller et al., 2011; Park et al., 2007; Lienkamp et al., 2012; Borovina et al., 2010), while canonical Wnt/β-catenin signaling is required for the specification and renewal of nephron progenitors (McNeill, H., 2009; Simons et al., 2005; Karner et al., 2009; Kishimoto et al., 2008). In addition to its influences on cell behaviors and morphology, including polarity establishment and division orientation (Lienkamp et al., 2012; Borovina et al., 2010), PCP signaling is also instrumental in facilitating the orientation of cell motility and coordinating cell interactions to facilitate nephric tubule formation (Saburi et al., 2008; Lienkamp et al., 2012; Miller et al., 2011). Epithelial nephron progenitors establish apical-basal polarity through the activity of the Par, Crumbs, and Scribble complexes, which define membrane domains via mutually antagonistic interactions and polarized trafficking (Goldstein and Macara, 2007; Rodriguez-Boulan and Macara, 2014). PCP further aligns epithelial cells across the tissue plane, coordinating convergent extension movements that drive tubule elongation (Karner et al., 2009; Saburi et al., 2008; Lienkamp et al., 2012). Initiation of PCP signaling results in the activation of the downstream formin protein Dishevelled-associated activator of morphogenesis 1 (Daam1) (Habas et al., 2001). Formin proteins facilitate actin polymerization by integrating profilin-associated monomers at the barbed, growing end of microfilaments. These polarity cues interface with the actomyosin cytoskeleton to regulate junctional maturation, rosette topology, and directional intercalation (Nishimura et al., 2012; Walck-Shannon and Hardin, 2014). In parallel, cadherin-based adherens junctions (AJs) form the adhesive structure of epithelia, linking neighboring cells to the actin cytoskeleton and thereby providing the mechanical framework that stabilizes epithelial architecture and tissue integrity (Harris & Tepass, 2010; Niessen et al., 2011; Takeichi, 2014). AJ assembly requires not only cadherin homophilic binding but also dynamic reinforcement through β-catenin, p120-catenin, and actin-associated adaptors, all of which depend on continuous cytoskeletal remodeling (Takeichi, 2014; Harris and Tepass, 2010; Yonemura et al., 2010).

A critical but often underappreciated component of junctional regulation is vesicular transport. Junctional proteins, including E-cadherin, undergo constant cycles of endocytosis, sorting, and recycling to maintain homeostasis and support tissue remodeling (Le et al., 1999; Bryant and Stow, 2004; Kowalczyk and Nanes, 2012). Endocytic pathways internalize junctional components during migration, damage, or morphogenetic rearrangements, while exocytic pathways deliver newly synthesized or recycled cadherins back to the plasma membrane (Harris and Tepass, 2010). Rab11-dependent recycling, the exocyst complex, and Cdc42-mediated vesicle docking are essential for polarized E-cadherin delivery, lumen formation, and epithelial integrity (Langevin et al., 2005; Lock and Stow, 2005; Bryant et al., 2010). Disruption of these pathways leads to intracellular cadherin accumulation, weakened junctions, and defective tubulogenesis (Bryant et al., 2014). These observations underscore that vesicular transport is not merely supportive but is fundamentally required for the assembly, maintenance, and remodeling of adhesive contacts in dynamic tissues (Bryant and Mostov, 2008; Rodriguez-Boulan and Macara, 2014). Despite the clear requirement for both vesicular transport and cytoskeletal regulation in junctional assembly, the molecular mechanisms that coordinate these systems during nephron development remain unresolved. However, Dynamin Binding Protein (Dnmbp) is a compelling molecular bridge candidate (Bryant and Mostov, 2008; Rodriguez-Boulan and Macara, 2014, Salazar et al., 2003).

Dynamin Binding Protein (Dnmbp) is a multidomain scaffolding protein involved in various cellular processes and regulatory pathways. The unique domain architecture of Dnmbp consists of four N-terminally located Src Homology-3 (SH3) domains, a centrally located Dbl Homology (DH) domain, and a C-terminally located BIN/amphiphysin/Rvs (BAR) domain that is followed by two additional SH3 domains (**Fig.1A**) The four N-terminal SH3 domains can independently bind dynamin, which is known for its role in lipid bilayer bending and scission during endocytosis (Salazar et al., 2003) as well as stimulate N- WASP-dependent actin assembly via its C-terminal SH3 domain (Kovacs et. al., 2006), resulting in a functional scaffold between endocytic and actin regulatory pathways (Salazar et al., 2003). In addition to the N-terminal binding capability of Dnmbp directly to dynamin, the C-terminal BAR domain of Dnmbp could also provide aid in dynamin scission as it has its own membrane bending capabilities (Carman & Dominguez, 2018). However, the N-terminal SH3 domains upstream of the Rho GTPase binding DH domain result in autoinhibition, blocking Cdc42 binding and activation, as well as membrane binding capability of the BAR domain (Carman & Dominguez, 2018). Prior studies, conducted in HEK293 cells have shown that the tricellular tight-junction-associated protein component tricellulin (a member of the Marvel protein family) releases the N-terminal SH3 intramolecular inhibition when the N-terminal domain of tricellulin binds to the SH3-C2 (SH3 number six) domain of Dnmbp (Oda et. al., 2014). An alternate model proposed to explain the release of the autoinhibiting N-terminal SH3 domain, speculates that protein truncation and/or post-translational modifications, including phosphorylation, could be a mechanism for removing the N-terminal regulator of DH containing proteins (Carman & Dominguez, 2018). Nevertheless, interaction between Dnmbp and Cdc42 continues to occur resulting in the regulation of Cdc42 GTPase activation. Previous work has shown that Dnmbp influences ciliary assembly and tubulogenesis through activation of Cdc42 (Baek et. al., 2016, Martin-Belmonte et al., 2007, Zuo et al., 2011), as well as being required for facilitating proper spindle orientation (Qin et. al., 2010) and junctional configuration in epithelial tissues (Otani et. al., 2006). As a key regulator of Cdc42, Dnmbp plays an essential role in vesicle transport, the establishment of cell polarity, tissue morphogenesis, and cell-cell junction assembly (Martin-Belmonte et al., 2007, Bryant et al., 2010; Baek et. al., 2016; Zuo et al., 2011). Disruption of Cdc42, which is activated via its interaction with the DH domain of Dnmbp, results in ciliary defects and cyst formation within the kidney (Zuo et al., 2011). Additionally, published work from the Miller lab has revealed that Dnmbp is essential for *Xenopus* embryonic kidney development and depletion of Dnmbp results in the disruption of nephrogenesis (DeLay et al., 2019). However, the mechanisms by which Dnmbp is influencing nephron development have yet to be elucidated.

**Figure 1.**
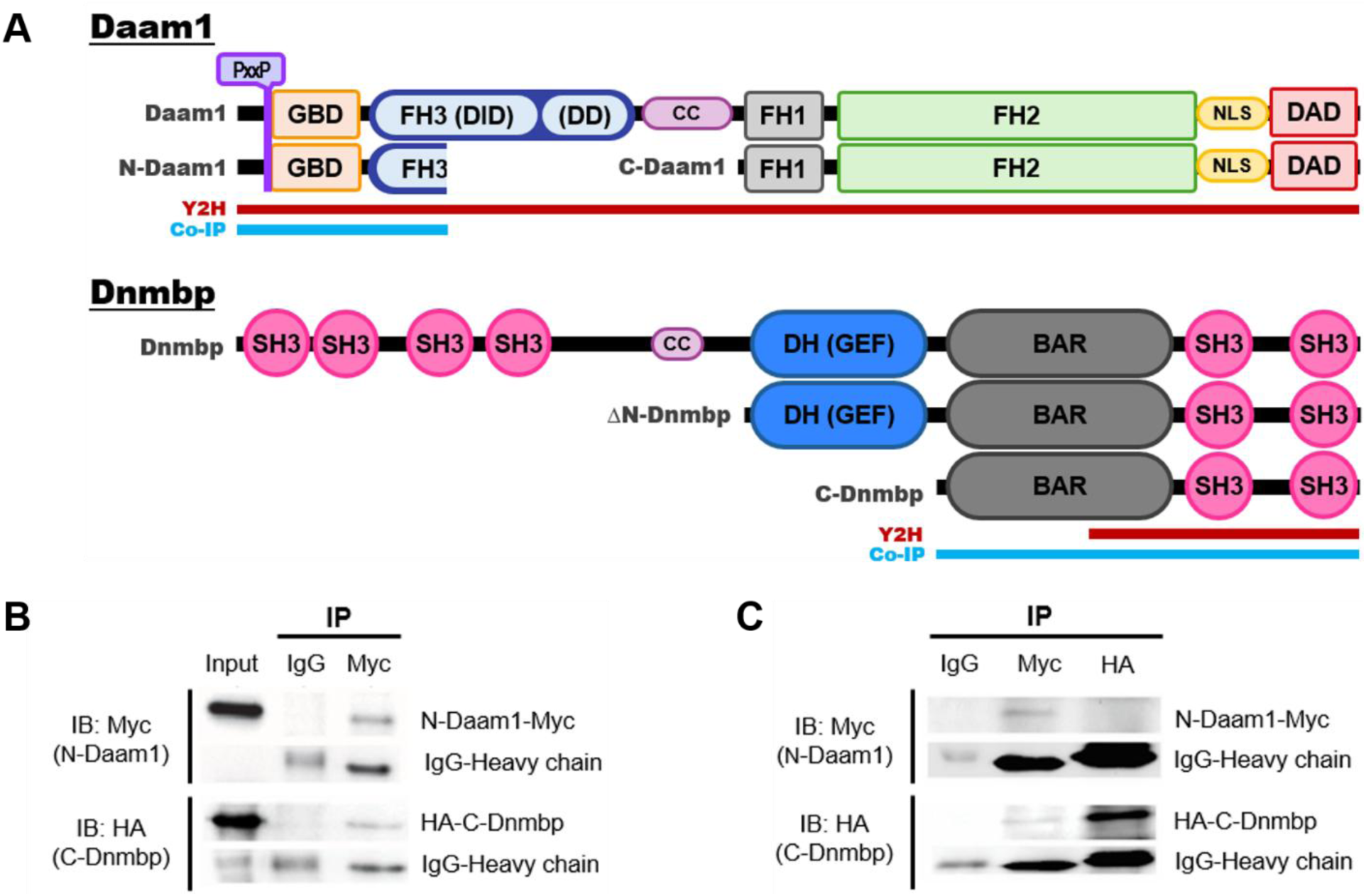
The N-terminus of Daam1 interacts with the C-terminus end of Dnmbp. A) Schematic of Daam1 and Dnmbp expression constructs. Red bars indicate interacting regions determined by the yeast two-hybrid (Y2H) screen. Blue bars indicate interacting regions based upon B) in vitro transcription and translation Co-immunopreciptiated (IP) experiments and C) exogenous mRNA in vivo translation Co-IP experiments. Immunoblots (IB) using Myc (top) and HA (bottom) antibodies show that upon IP of N-Daam1-Myc with a Myc antibody, HA-C-Dnmbp co-IPs. Note: Domains are not to scale. GBD = GTPase-binding domain; FH = formin homology; CC = coiled coil; NLS = nuclear localization sequence; DAD = diaphanous autoinhibitory domain; SH3 = Src homology 3; DH = Dbl homology; BAR = BIN/amphiphysin/Rvs

Here, we identify Dnmbp as a novel interacting partner of the Wnt/PCP formin Daam1 and define its role during nephron development. Our findings demonstrate that Dnmbp is required for proper assembly of E-cadherin-containing cell-cell junctions, revealing a previously unrecognized mechanism linking Wnt/PCP signaling to epithelial adhesion during nephrogenesis.

## RESULTS

### Dnmbp interacts with Daam1 within nephric primordium

To identify potential ligands for the Wnt/PCP formin protein Daam1 during nephrogenesis, a comprehensive yeast two-hybrid (Y2H) screen was conducted using an embryonic mouse kidney library. Dnmbp emerged as a recurrent candidate in two independent screens encompassing approximately 150 million yeast clones. Using full-length Daam1 as the “bait”, the screens revealed an interaction with the C-terminal region of Dnmbp, containing the distal end of the BAR domain and two Src Homology 3 (SH3) domains (**Fig. 1A**, red bars). SH3 domains are well-established mediators of protein-protein interactions and frequently engage with regulators of the actin cytoskeleton (Mayer B. J., 2001; Cestra et al., 2005; Kovacs et. al., 2006), supporting the possibility of a functional interaction between Dnmbp and Daam1. To validate the interaction identified in the Y2H assay, an HA-tagged Dnmbp C-terminal construct (C-Dnmbp) containing the domains recovered in the screen was generated (**Fig. 1A**). Protein expression was confirmed by both an *in vitro* transcription and translation (TnT) assay (**Fig. 1B**) and an exogenous mRNA translation assay (**Fig. 1C**). Co-immunoprecipitation (Co-IP) analyses demonstrated that the C-terminus of Dnmbp, which includes a classic BIN/amphiphysin/Rvs (BAR) domain and two SH3 domains, interacts with the N-terminal region of Daam1 (N-Daam1) containing the GTPase binding domain (GBD) (**Fig. 1**, blue bars). Previous studies have reported that the GBD domain of Daam1 interacts with both active (GTP-bound) and inactive (GDP-bound) states of Cdc42, although it has a higher avidity for GTP-Cdc42 (Aspenström et al., 2006; Liu et al., 2008). The centrally located DH domain of Dnmbp is a known activator of GDP-Cdc42 (Urrutia et. al., 2021; Cestra et al., 2005; Kovacs et. al., 2006; Kovacs et. al., 2011; Bruurs et al., 2018). However, for Dnmbp to exhibit guanine nucleotide exchange factor (GEF) activity of the DH domain, it must be stimulated to release autoinhibitory interactions that occur between the N- and C-terminus that sequester the DH domain (Salazar et al., 2003; Oda et al., 2014). Proline-rich sequences often serve as recognition motifs for SH3 domains, especially PxxP motifs (where P= proline and x= any amino acid) (Kay et al., 2000). These PxxP motifs have been shown to interact with the final C-terminus SH3 domain (SH3-C2) of Dnmbp to release its autoinhibition, enabling GEF activity (Cestra et al., 2005; Kovacs et. al., 2011; Bruurs et al., 2018; Oda et al., 2014). Collectively, these findings suggest that interaction of Daam1 with the SH3-containing C-terminus of Dnmbp may represent a potential mechanism for coupling Daam1 signaling to Dnmbp-mediated regulation of Cdc42 activity and actin cytoskeletal remodeling.

### Structural modeling predicts an interaction between Daam1 and Dnmbp

To further examine the molecular basis of the interaction identified by yeast two-hybrid screening and confirmed by co-immunoprecipitation, we generated a structural model of the Daam1-Dnmbp complex using AlphaFold3 (**Fig. 2**).

**Figure 2.**
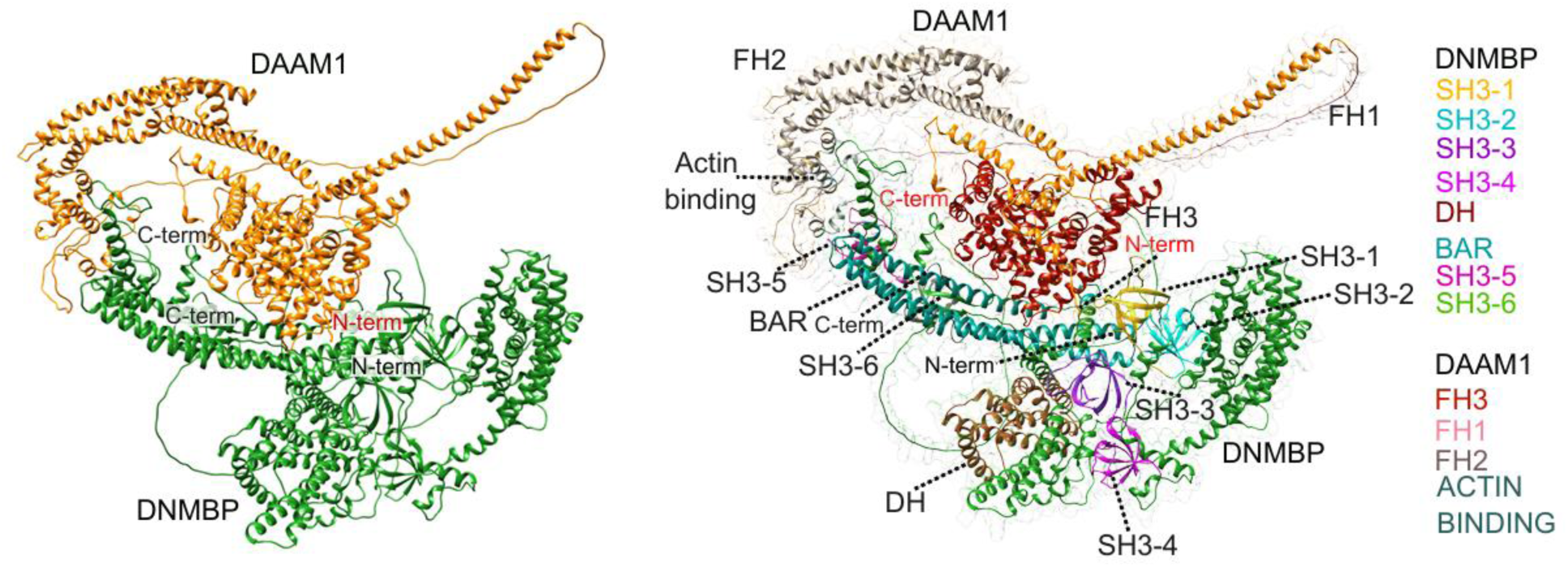
AlphaFold3 structural model predicts an interaction between the N-terminal region of Daam1 and the C-terminal SH3/BAR-containing region of Dnmbp. Structural model of the Daam1–Dnmbp complex generated using AlphaFold3. Dnmbp contains six Src homology 3 (SH3) domains, designated SH3-1 (amino acids 2–61; yellow), SH3-2 (66–126; cyan), SH3-3 (145–204; purple), SH3-4 (243–302; magenta), SH3-5 (1285–1348; violet red), and SH3-6 (1513–1576; light green), together with the Dbl homology (DH) domain (784–967; sienna) and the BIN/amphiphysin/Rvs (BAR) domain (1008–1217; dark cyan). Daam1 consists of the FH3 domain (45–420; dark red), the proline-rich FH1 domain (528–599; plum), and the FH2 domain (600–1009; dark gray), which contains the actin-binding region (693–702; light blue). FH = formin homology

The predicted complex places the N-terminal FH3 domain of Daam1 in close proximity to the BAR domain and the N-terminal SH3 domains of Dnmbp, while the C-terminal domains, SH3-5 and SH3-6, are positioned far from proximity from the primary interaction interface. This configuration aligns with biochemical investigations indicating that the N-terminal region of Daam1 interacts with the C-terminal area of Dnmbp, as identified in the yeast two-hybrid screen. Instead of showing a single domain-domain interaction, the model presents an extensive interaction interface that encompasses both the BAR domain and neighboring SH3 domains.

### Dnmbp localization during embryonic kidney development

As a first step to elucidate the role Dnmbp plays during nephron epithelialization and visualize the subcellular localization patterns of Dnmbp *in vivo*, fluorescently labeled DNA constructs were engineered of full-length human DNMBP (hDNMBP) and a constitutively active C-terminal variant devoid of the four N-terminal SH3 domains, referred to as ΔN-hDNMBP. The N-terminal SH3 domains were specifically targeted based on Dnmbp recruitment of actin-regulating proteins via the C-Terminal SH3 domains in neuronal cells (Salazar, 2003; Kovacs et al., 2006; Otani et al., 2006). Both constructs encompass the region of Dnmbp indicated in the yeast two-hybrid screen (**Fig. 1**, red bars) to be responsible for interacting with Daam1. Using a transgenic Xla.Tg(pax8:GFP) *Xenopus* kidney reporter line, subcellular localization of each protein was evaluated in fixed NF stage 28 (the time in which epithelialization is beginning) embryonic kidneys during early nephrogenesis. Consistent with the prior observations seen in the epidermis (data not shown), hDNMBP (**Fig.3A**) localized in scattered puncta throughout the cell, whereas the constitutively active ΔN-hDNMBP was highly cytoplasmic (**Fig.3B**). At this stage, in these embryos, defined nephric cell-cell junction membranes were not yet strongly evident, however a few (presumably newly forming) cell-cell borders with minimal intensities were noted.

**Figure 3.**
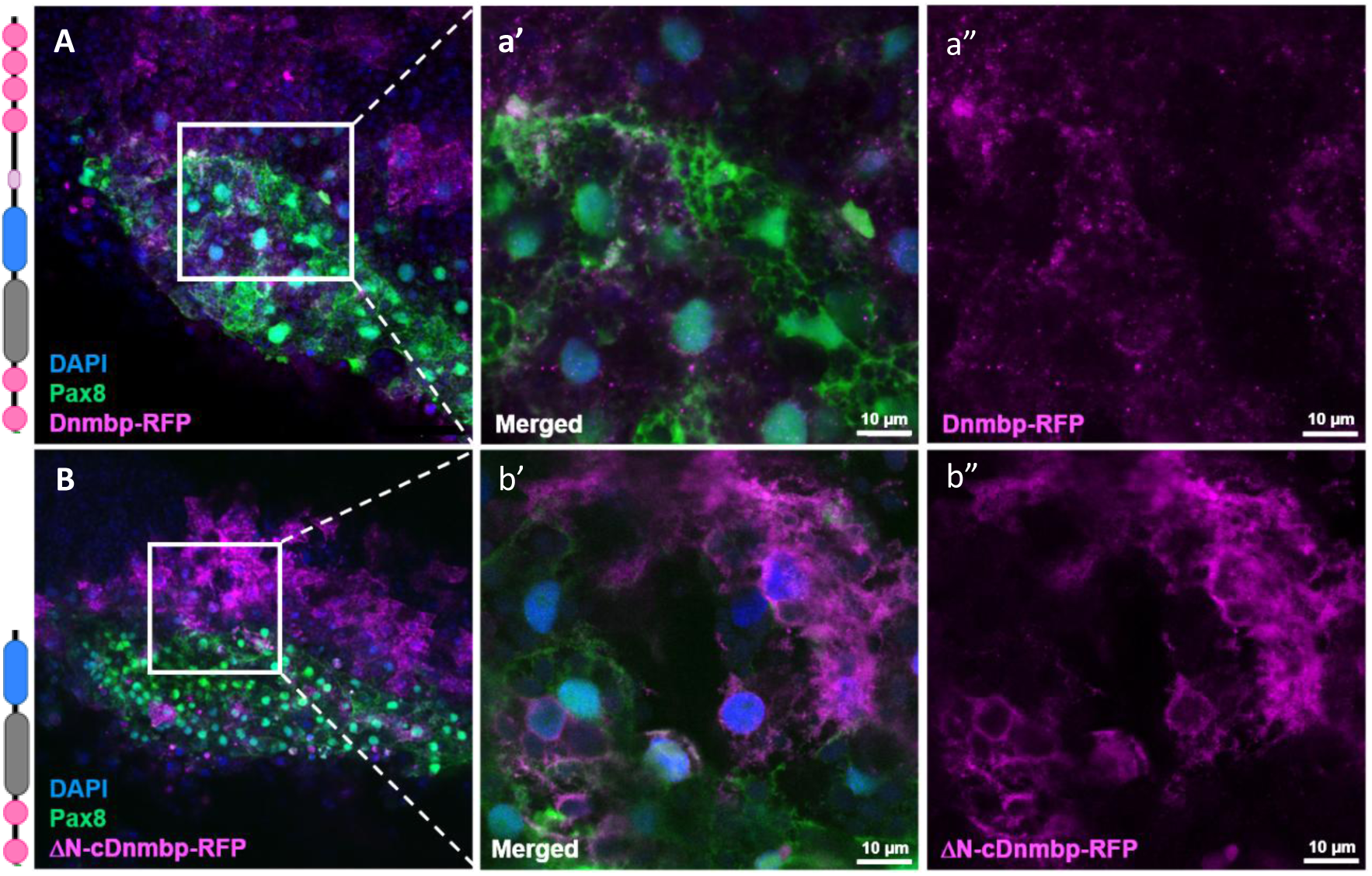
Overexpression of DNMBP in *Xenopus* embryonic kidney cells. Confocal maximum image projections of whole-mount immunostaining of *Xenopus* nephron progenitor cells, marked by transgenic *pax8::gfp* (green) reporter expression. Top panel (A) Full-length hDNMBP-RFP (magenta) construct localization within nephric primordium; Bottom panel (B) ΔN-hDNMBP-RFP (magenta) lacking N-terminal SH3 domains within nephric primordium; a’-a” and b’-b” show close-up views of white boxes. Scale bars measure 10 µm.

Therefore, to assess subcellular hDNMBP localization in epithelialized tissue, 0.5 ng of either fluorescently tagged full length *hDNMBP* mRNA or the constitutively active *ΔN-hDNMBP* mRNA were injected into the V2 blastomere of 8-cell stage *Xenopus* embryos to target the nephric primordium. Subcellular localization of the fluorescent proteins was detected by whole-mount immunofluorescent staining of stage 40 embryos fixed in methanol-free fixative prior to staining with antibodies against GFP or RFP to visualize the hDNMBP tag, against β-catenin, a structural component fundamental to cadherin-based adherens junctions, and against 3G8 and 4A6 to label mature nephrons as well as phalloidin staining to visualize F-actin. Upon imaging the mature nephron, the full length hDNMBP protein appeared to have strikingly different localization patterns compared to the constitutively active protein lacking the four N-terminal SH3 domains (**Fig. 4**).

**Figure 4.**
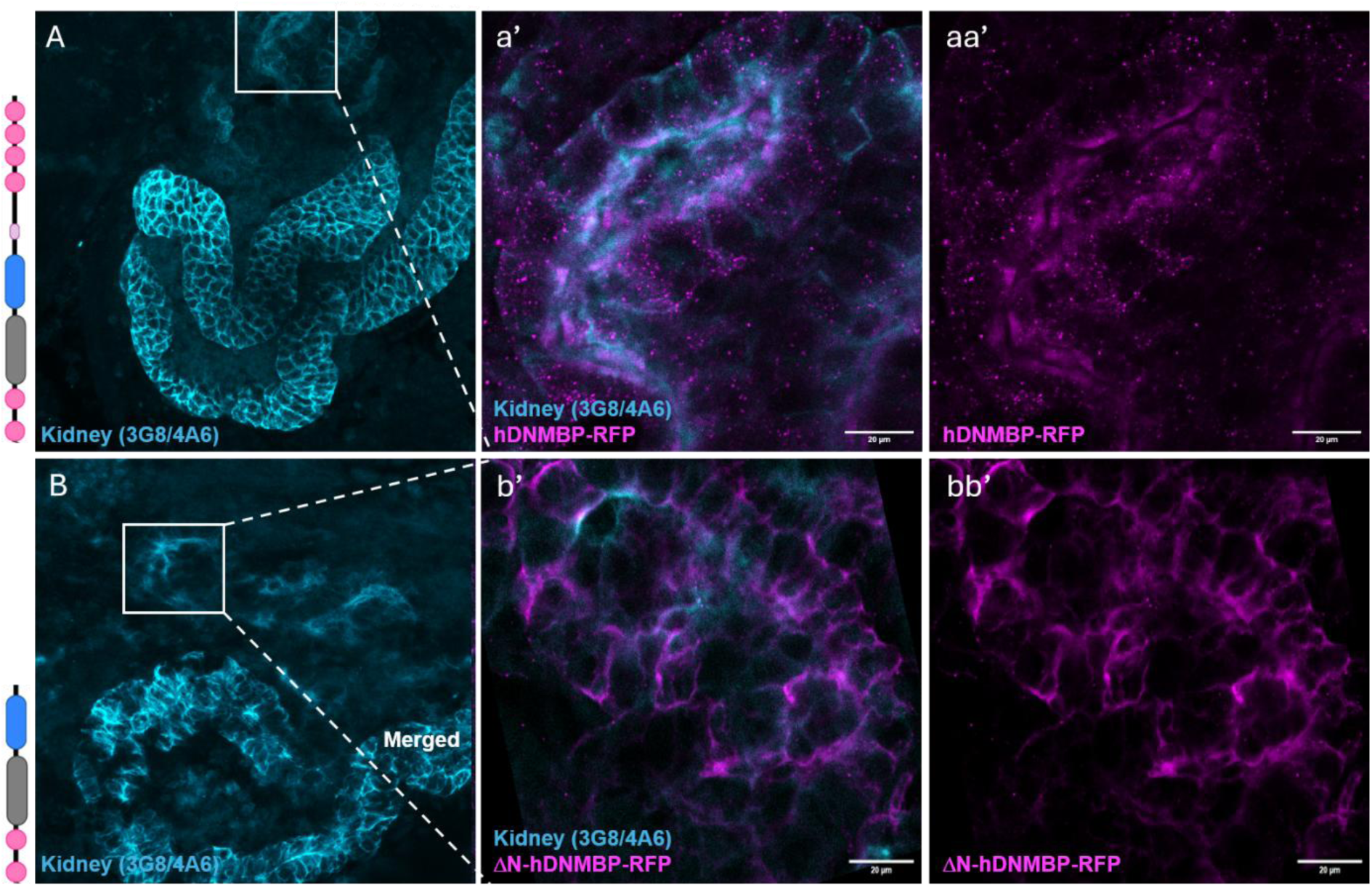
Overexpression of *DNMBP* in mature *Xenopus* pronephros. Confocal maximum image projections of mature *Xenopus* pronephros, marked by 3G8 for proximal tube lumen and 4A6 for distal duct membranes (blue). (A) Exogenous full-length hDNMBP-RFP (magenta) localizes predominantly at puncta within nephric tubules; (B) Constitutively active ΔN-hDNMBP-RFP (magenta) lacking N-terminal SH3 domains localizes primarily to cell membranes in *Xenopus* nephric epithelia; a’-a” and b’-b” show close-up views of white boxes. Scale bars measure 20 µm.

To determine whether the observed scattered puncta of hDNMBP were localizing to cell-cell contacts in an unobvious manner, four-cell embryos were co-injected with mRNAs to encode the full-length hDNMBP containing a RFP tag (pCS2-hDNMBP-RFP) and a GFP tagged E-cadherin (pCS2-Ecad-GFP) construct. Embryos were first observed under high-resolution confocal microscopy at approximately NF stage 28; however, GFP tag intensity was minimal (data not shown). Therefore, the embryos were left to continue developing and were observed again when epithelialization was well underway (NF stage 34/36). Upon re-evaluation, dynamic interactions between the scattered puncta of exogenous full-length hDNMBP were observed closely associating with the tag-GFP of E-cadherin at cell-cell contacts. (**Fig.5**). In addition to a strong presence at cell membrane contacts, E-cadherin-GFP was localized around spherical structures aggregating to what are presumed to be adhering bicellular, tricellular, and rosette-like cell-cell junctions. hDNMBP-RFP expression was visible in association with, or in close proximity to, many of the circular structures, including several instances in which hDNMBP fluorescent expression could be seen filling the E-cadherin sphere. However, due to the toxicity of E-cadherin overexpression, only one surviving embryo could be evaluated.

**Figure 5.**
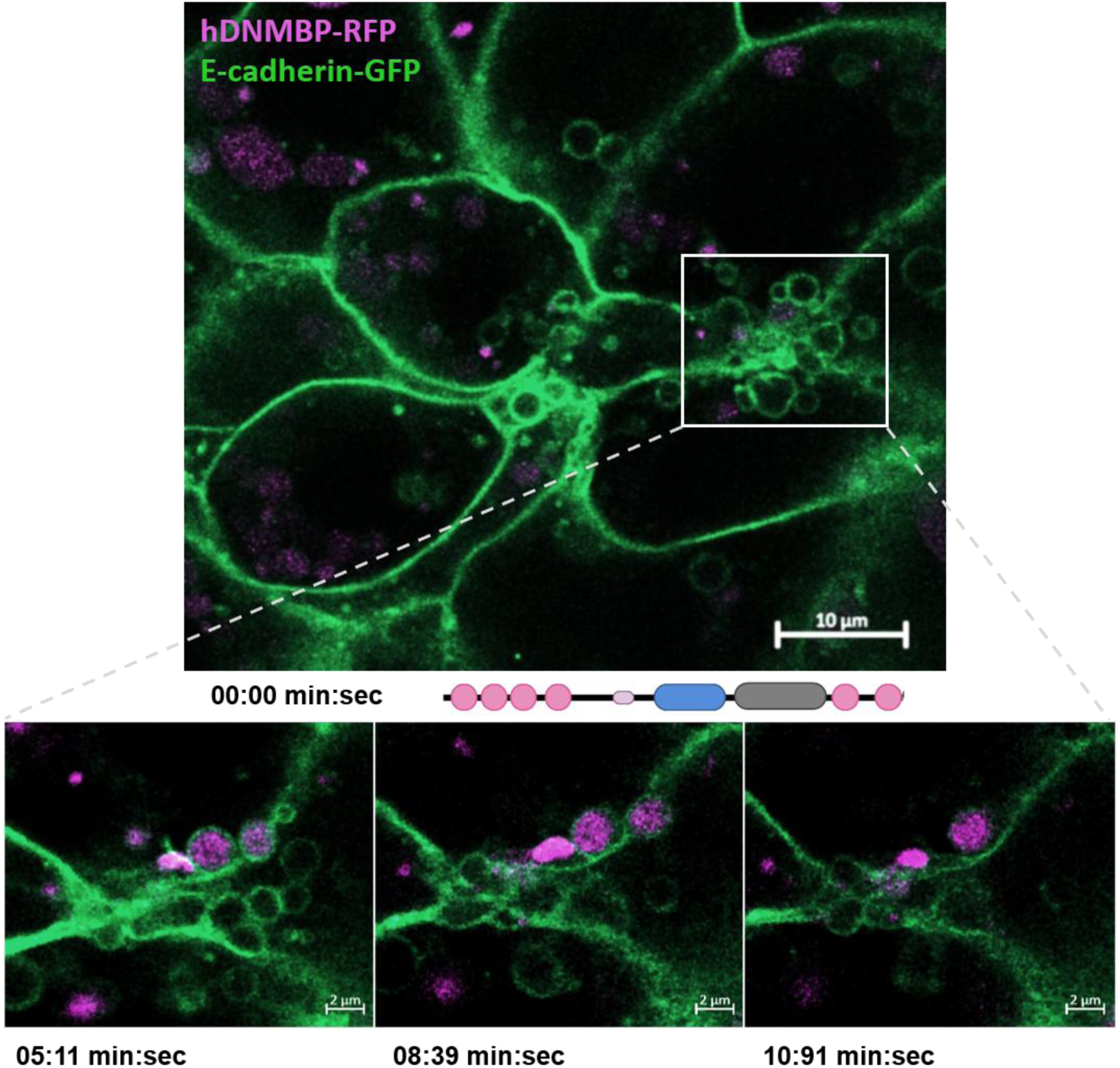
Dnmbp associates with E-cadherin vesicle-like structures in *Xenopus* kidney. Live super-resolution time-lapse imaging in *Xenopus* windowed kidneys shows hDNMBP (magenta) and E-cadherin (green) constructs localizing in close association at cell-cell junctions in epithelializing nephric primordium. Bottom three panels represent close-up images of the white box in the top panel at the indicated times below.

### Dnmbp depletion disrupts E-cadherin localization in developing nephrons

To assess the influence of Dnmbp on adherens junction formation during nephron epithelialization, E-cadherin localization at cell-cell junctions was observed upon Dnmbp morpholino-mediated inhibition. To predict the effect of Dnmbp-depletion upon junctional integrity, several points were considered: 1. Morpholino (MO)-mediated inhibition within the target tissue often results in mosaic knockdown. 2. Dnmbp-MO1 only blocks translation of the full-length Dnmbp (isoform-a) protein because other protein isoforms are encoded by mRNAs contain different 5’UTRs. Therefore, not all endogenously expressed Dnmbp in the targeted nephric primordium was depleted.

The data collected over four independent trials show that E-cadherin fails to localize to the membranes in epithelializing nephrons of *Xenopus* tadpoles upon Dnmbp depletion (**Fig. 6A**). Acquired fluorescence imaging suggests lateral diffusion (data not shown) and reduced localization of E-cadherin to the cell-cell membranes of Dnmbp knockdown (KD) samples, as well as undulating and/or broken membrane borders. Relative intensity profiles of E-cadherin fluorescence at cell-cell contacts were analyzed by one-way ANOVA, followed by *Tukey HSD,* which showed that std-Control vs Dnmbp KD was significantly below the threshold with an adjusted p-value of P < 0.0001, as was the iControl KD vs Dnmbp KD with an adjusted p-value of P < 0.0001 (**Fig. 6B**). However, when comparing the two control samples (std-Control vs iControl KD) the adjusted p-value was calculated at P = 0.9970 and did not fall below the dictated threshold. Additionally, data from Western blot analysis indicated the total E-cadherin concentration in Dnmbp-depleted samples is not reduced (**Fig. 6C**).

**Figure 6.**
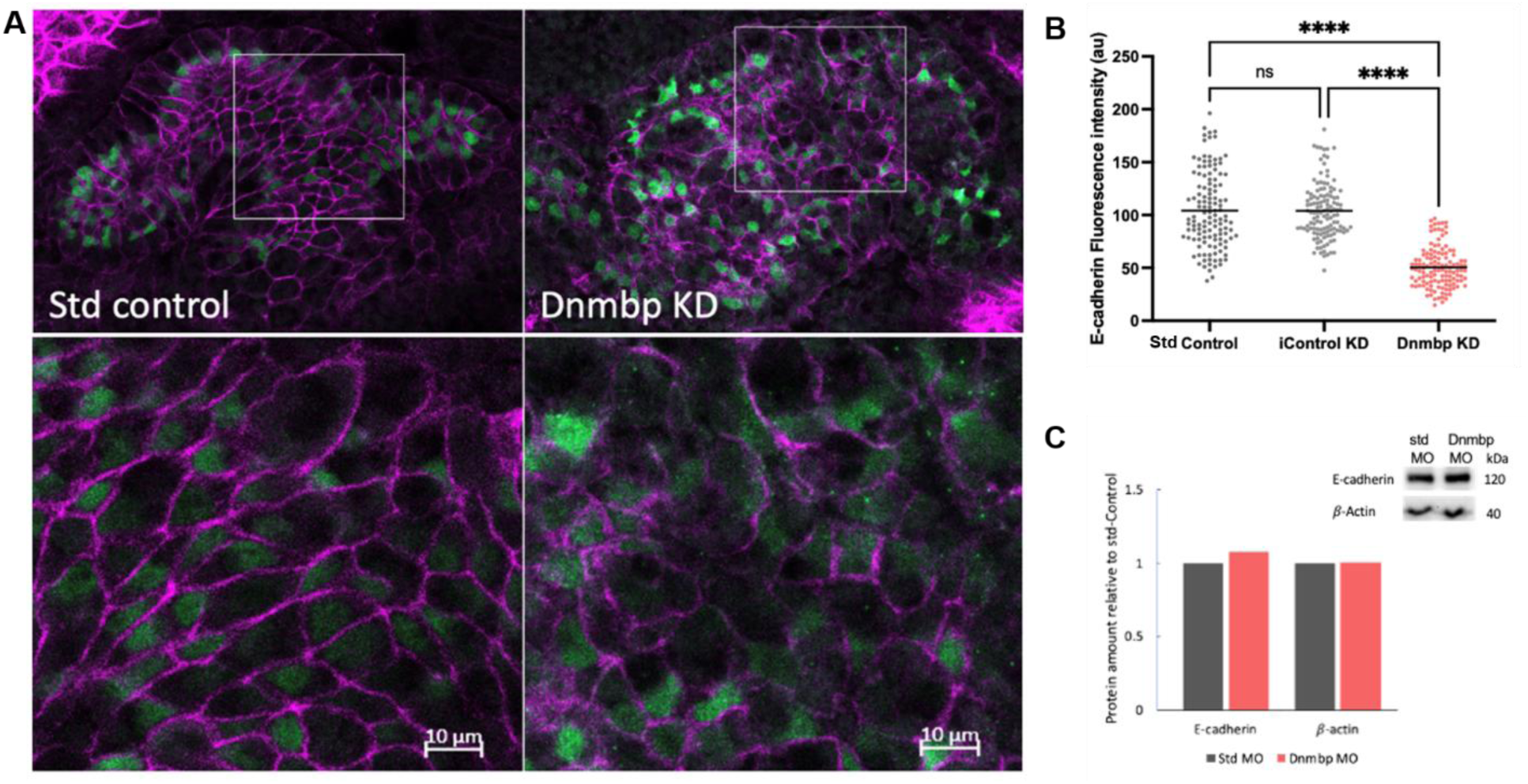
Dnmbp is required for E-cadherin localization to the junctions in epithelializing nephrons. A.) Maximum projection confocal images of E-cadherin (magenta) expression (top) within epithelializing nephric primordium (marked by Lhx1 antibodies; green) in Std Control and Dnmbp KD embryos (confocal image of iControl KD not shown). Bottom panels represent white box regions of interest. Scale bars measure 10 µm. B.) Scatterplot showing the relative distribution of E-cadherin fluorescence intensity in nephron progenitors of std-control, iControl KD and Dnmbp KD embryos. N_Std-Control_=126 junctions over 4 trials, N_iControl_=126 junctions over 4 trials, and N_Dnmbp KD_=126 junctions over 4 trials. ****P < 0.0001 analyzed by one-way ANOVA. C.) Western blot and corresponding graph of densitometry for E-cadherin and β-actin (control) protein levels for Std Control (Std MO) and Dnmbp KD (Dnmbp MO) injected embryos.

To distinguish between persistent disruption and delayed recruitment of junctional E-cadherin, its localization at mature nephron cell junctions was analyzed following Dnmbp depletion. Stage 40-42 embryos fixed by formalin fixation were stained with primary antibodies against RFP to confirm the presence of co-injected membrane-RFP tracer and against E-cadherin to assess junction integrity. Dylight-594 tomato Lectin was used for mature nephron labeling, as well as DAPI staining to label nuclei. Quantification of junctional integrity was achieved by measuring the fluorescent intensity of E-cadherin along individual pronephric cell-cell borders with confirmed RFP presence in both adjacent cells (+RFP). Analysis of E-cadherin intensity in the Dnmbp KD kidney was compared with that of the iControl and Std Control kidneys using Prism software. The data indicated that depletion of Dnmbp within the mature *Xenopus* embryonic nephron significantly disrupts the localization of junctional E-cadherin (**Fig. 7**). To address whether RFP negative cells adjacent to RFP positive cells have E-cadherin defects, fluorescent intensity profiling of E-cadherin along individual cell-cell borders when only one cell had RFP expression present, as well as along cell-cell contacts when neither cell had RFP expression present (Dnmbp KD-RFP). In addition, fluorescent intensity profiling of junctional E-cadherin in mature nephron cells of the iControl kidney (image not shown) was also quantified. Analysis of E-cadherin intensity in the Dnmbp KD kidney compared with the Std Control kidney was performed using Prism software. The mean fluorescence intensities used for analysis of iControl cell junctions were acquired from the same embryos in which the injected side data were collected, and the same number of junctions were profiled for each side within each embryo in the trial. Data analysis indicates that E-cadherin localization to the junctions is significantly disrupted in the Dnmbp-depleted mature nephron, even when RFP tracer is not present in the junctions of injected kidney nephron progenitor cells (**Fig. 7**). The data also indicate that junctional E-cadherin localization of iControl kidneys is not significantly affected by Dnmbp depletion treatments on the other side of the embryo.

**Figure 7.**
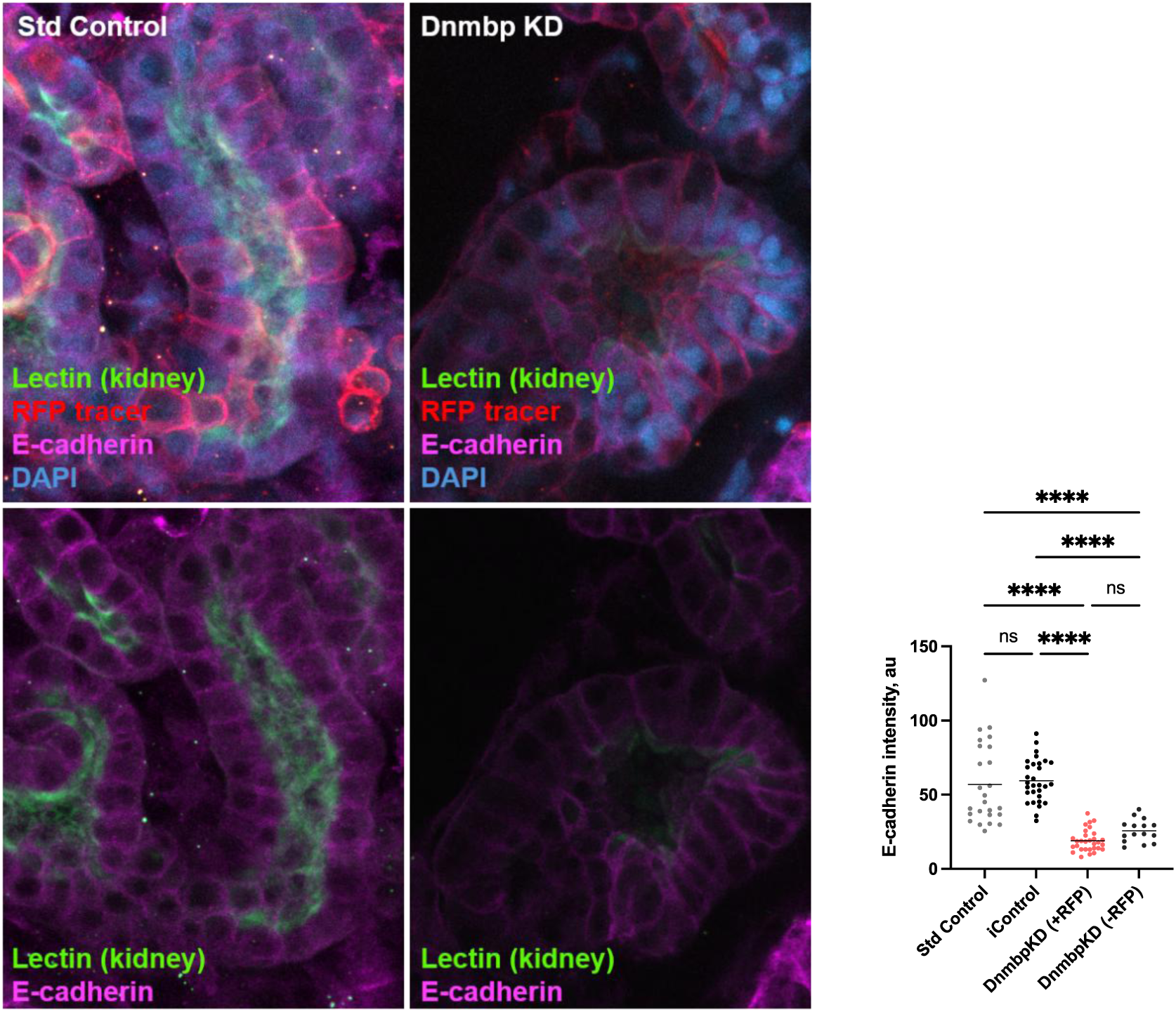
Dnmbp is required for E-cadherin localization to the junctions in mature nephrons. Dnmbp is required for E-cadherin localization to the junctions in mature nephrons. Maximum projection confocal images of E-cadherin (magenta) localization in mature *Xenopus* proximal tubule (marked by Dylight-488 Lectin), RFP injection tracer (Red), and DAPI (Blue) nuclear stain in Std Control and Dnmbp KD embryos (confocal image of iControl not shown). Scatterplot shows the relative distribution of E-cadherin along individual pronephric cell-cell borders within the Dnmbp-KD kidney compared to the uninjected internal control (iControl) kidney and standard morpholino control (Std Control) kidney with confirmed RFP presence in both adjacent cells (+RFP). E-cadherin along individual cell-cell borders when only one cell had RFP expression present, as well as along cell-cell contacts when neither cell had RFP expression present (DnmbpKD-RFP), was also analyzed. ****P < 0.0001 analyzed by one-way ANOVA.

#### Human DNMBP rescues junctional E-cadherin upon depletion of *Xenopus* Dnmbp

To verify that Dnmbp loss of function was responsible for the reduction of E-cadherin at the junctions, kidney-targeted microinjection of mRNA encoding full-length (FL) hDNMBP was utilized to conduct loss-of-function rescue experiments (**Fig. 8**). As a negative mRNA control, *β*-*galactosidase* (*β-gal*) was co-injected with the standard control morpholino, as well as the Dnmbp morpholino. As expected, confocal imaging of E-cadherin within nephron progenitors indicated a significant reduction of junctional E-cadherin in the Dnmbp-MO1 and *β-gal* treatment (Dnmbp KD) when compared to the standard control and *β-gal* treatment (std control) (mean diff.= 223.1; P <0.0001). Co-injection of Dnmbp-MO1 and mRNA encoding full length hDNMBP (DNMBP rescue) resulted in a significant increase of junctional E-cadherin when compared to the Dnmbp KD (mean diff.= -177.9; P <0.0001), although not to wild-type levels. These data indicate that the loss of E-cadherin at cell-cell contacts is due specifically to Dnmbp depletion within the nephric primordium.

**Figure 8.**
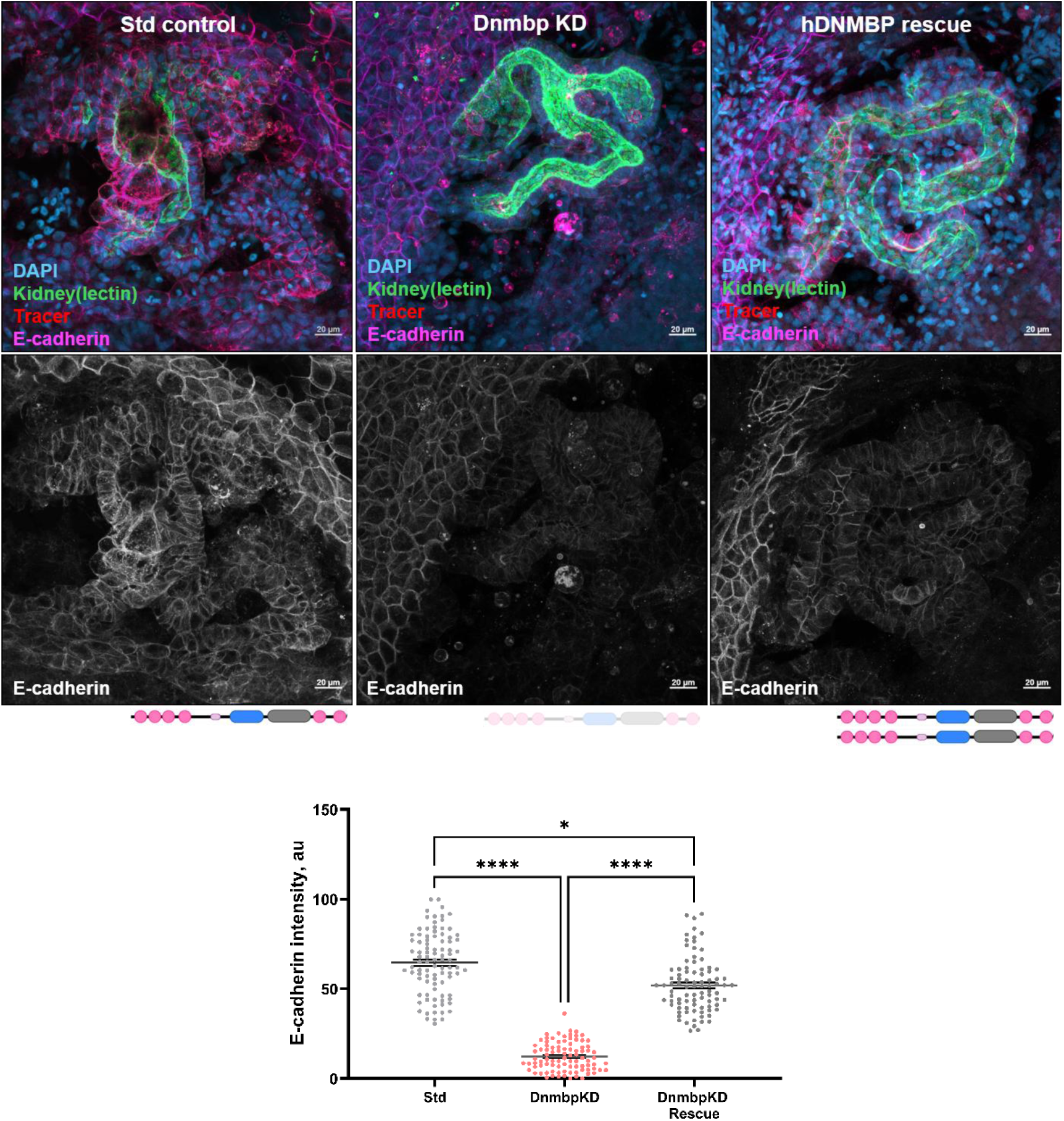
Full-length human DNMBP rescues E-cadherin localization to adherens junctions upon depletion of *Xenopus* Dnmbp. Confocal imaging of E-cadherin within mature nephrons (marked by Dylight-488 Lectin) indicates a significant reduction of junctional E-cadherin upon morpholino-mediated Dnmbp depletion when compared to standard control morpholino, FL-hDNMBP mRNA rescue of *Xenopus* Dnmbp KD. Scale bars measure 20 µm. Scatterplot shows relative intensity of junctional E-cadherin in mature epithelialized pronephros, quantified using GraphPad Prism Software. N_std-Control_=85 junctions over 3 trials, N_DnmbpKD_=85 junctions over 3 trials, and N_rescue_=85 junctions over 3 trials nsP > 0.05, *P ≤ 0.05, **P ≤ 0.01, ****P ≤ 0.0001 analyzed by one-way ANOVA

## DISCUSSION

Nephron formation requires the precise coordination of cell polarity, cell-cell adhesion, cytoskeletal remodeling, and membrane trafficking as mesenchymal progenitors undergo epithelialization and organize into a functional tubular epithelium (Costantini and Kopan, 2010; Little and McMahon, 2012; Dressler, 2006; Bryant and Mostov, 2008). Here, we identify Dnmbp as a novel interacting partner of Daam1 and provide evidence that Dnmbp is required for the localization of E-cadherin to nephron cell-cell contact sites during epithelialization. Together, our findings support a model in which Dnmbp and Daam1 cooperate to coordinate cytoskeletal organization, Cdc42 signaling, and adherens junction maturation during nephrogenesis.

The identification of Dnmbp in two independent Y2H screens strongly suggests that the interaction between Daam1 and Dnmbp is biologically relevant. Subsequent Co-IP experiments confirmed that the SH3-containing C-terminal region of Dnmbp interacts with the N-terminal region of Daam1 containing the GTPase-binding domain. This interaction is particularly intriguing given that Dnmbp functions as a Cdc42-specific guanine nucleotide exchange factor whose activity is normally constrained by autoinhibitory interactions involving its C-terminal SH3 domains (Salazar et al., 2003; Cestra et al., 2005; Oda et al., 2014; Bruurs et al., 2018). Daam1 functions as a formin downstream of non-canonical Wnt signaling that promotes actin polymerization and cellular morphogenesis (Habas et al., 2001), whereas Dnmbp is known as a Cdc42-specific guanine nucleotide exchange factor involved in epithelial polarity, membrane trafficking, and junctional organization (Cestra et al., 2005; Bruurs et al., 2018; Baek et al., 2016; Qin et al., 2010; Otani et al., 2006). Previous studies have demonstrated that Dnmbp activity is autoinhibited through intramolecular interactions involving its C-terminal SH3 domains and that binding of proline-rich ligands can relieve this inhibition and activate Cdc42 signaling (Oda et al., 2014; Bruurs et al., 2018). The recognized function of SH3 domains in facilitating protein-protein interactions and forming signaling complexes involved in actin cytoskeletal control suggests that the predicted structural modeling complex provides a credible structural foundation for the Daam1–Dnmbp interaction. One possibility is that Daam1 associates with the C-terminal region of Dnmbp to regulate Dnmbp activation, thereby linking Wnt/PCP signaling to localized Cdc42 activation. Although the model does not directly address the conformational changes required for Dnmbp activation, the proximity of Daam1 to the regulatory SH3/BAR region is consistent with the proposed mechanism in which Daam1 binding may influence Dnmbp autoinhibition.

Dnmbp localization studies provide further support for its role in epithelial junction formation. During nephrogenesis, full-length hDNMBP displayed a punctate distribution throughout nephric cells, whereas constitutively active ΔN-hDNMBP lacking the N-terminal SH3 domains was diffusely cytoplasmic. The distinctively different subcellular localization patterns of these constructs suggest that intramolecular regulatory domains influence not only Dnmbp activity but also its subcellular targeting. As epithelialization progressed, hDNMBP puncta were closely associated with exogenous E-cadherin-rich cell-cell contacts and circular membrane structures. Although the toxicity associated with E-cadherin overexpression limits the number of embryos available for analysis, these observations suggest that Dnmbp may localize to sites of active adherens junction remodeling. Such a localization pattern is consistent with previous reports demonstrating enrichment of Dnmbp at epithelial junctions, where it regulates Cdc42 activity and organization of the junctional actin cytoskeleton (Otani et al., 2006; Qin et al., 2010).

Loss-of-function analysis revealed that Dnmbp is required for proper junctional E-cadherin localization during nephron development. Depletion of Dnmbp resulted in a significant reduction of E-cadherin fluorescence at cell-cell borders, accompanied by discontinuous and wavy membrane boundaries. Importantly, Western blot analysis showed that total E-cadherin levels were unchanged, indicating that Dnmbp primarily affects E-cadherin localization rather than overall protein expression. This distinction suggests defects in protein transport, membrane retention, clustering, or cytoskeletal stabilization rather than altered E-cadherin synthesis. Such a role is consistent with previous studies showing that Dnmbp regulates membrane trafficking pathways and epithelial membrane organization through its effects on Cdc42 signaling and actin remodeling (Martin-Belmonte et al., 2007; Cestra et al., 2005; Kovacs et al., 2006). Furthermore, the persistence of junctional defects in mature nephrons indicates that Dnmbp is not simply required for the initial recruitment of E-cadherin but contributes to the long-term maintenance and stabilization of adherens junctions. The observation that E-cadherin defects extended beyond cells containing detectable lineage tracer suggests that Dnmbp-dependent junctional defects may influence neighboring cells within the epithelial network. Because adherens junctions are cooperative structures requiring contributions from both adjacent cells, disruption of E-cadherin recruitment in one cell may compromise the stability of junctions shared with neighboring cells that retain normal Dnmbp expression. Such non-cell-autonomous effects have been observed in other models of epithelial adhesion and underscore the importance of coordinated junction assembly across tissues (Harris and Tepass, 2010; Niessen et al., 2011). Alternatively, the incomplete correlation between tracer expression and morpholino distribution could simply reflect the mosaic nature of kidney-targeted microinjections. Regardless, the maintenance of normal E-cadherin localization within uninjected internal control kidneys argues against a systemic developmental defect and supports a local requirement for Dnmbp within the nephric primordium.

The successful rescue of junctional E-cadherin localization by full-length human *DNMBP* mRNA confirms the specificity of the morpholino and demonstrates functional conservation between *Xenopus* and human *DNMBP*. Although rescue did not fully restore E-cadherin to wild-type levels, the significant recovery of junctional localization strongly supports the conclusion that Dnmbp loss of function directly underlies the observed adhesion defects. Incomplete rescue could reflect differences in expression levels, timing of exogenous mRNA translation, or the inability of overexpressed protein to fully recapitulate endogenous spatial regulation.

Collectively, our data support a model in which Dnmbp cooperates with Daam1 to regulate the assembly and stabilization of nephron adherens junctions. We propose that interaction between Daam1 and the C-terminus of Dnmbp promotes localized Cdc42 activation at sites of epithelial remodeling. Activated Cdc42 could then coordinate Par-complex signaling, actin cytoskeletal organization, and membrane transport pathways required for efficient recruitment and stabilization of E-cadherin at developing junctions (Harris and Tepass, 2010; Georgio et al., 2008). Simultaneously, Daam1-mediated actin polymerization would provide the cytoskeletal framework necessary to support E-cadherin clustering and junction maturation (Krneta-Stankic et al., 2021). Therefore, disruption of the Dnmbp-Daam1 interaction would be expected to disrupt cytoskeletal remodeling during junction assembly, leading to reduced junctional E-cadherin, weakened epithelial architecture, and impaired nephron morphogenesis. This model provides a mechanistic link between Wnt/PCP signaling, Cdc42 activation, and adherens junction formation and identifies Dnmbp as a previously unrecognized regulator of kidney development. Future studies examining the effects of Dnmbp depletion on Daam1 localization, Cdc42 activity, and vesicular transport will further define the molecular mechanisms through which this signaling complex regulates nephrogenesis.

## ACKNOWLEDGMENTS

We thank Miller lab members as well as Dr. Pierre McCrea, Dr. Jae-II Park, Dr. Malgozata Kloc, and their lab members for lively discussions and suggestions on the manuscript. We also thank Dr. Yoshihiro Komatsu, Dr. George Eisenhoffer, Dr. Jeffrey Frost, Dr. Ruth Heidelberger, Dr. Guangwei Du, and Dr. Adriana Paulucci for their input and suggestions on the project. We are also thankful to Raymond Habas, Bruce Goode, Norihiro Sudou, and Masanori Taira for providing antibodies and constructs. We thank the instructors and teaching assistants of the 2023 Cold Spring Harbor Laboratory Cell & Developmental Biology of *Xenopus*: Gene Discovery & Disease Course. We are grateful to Jamie Cochran and Thomas H. Gomez, who take care of the animals. We are grateful to the UTHealth Office of the Executive Vice President and Chief Academic Officer and the Department of Pediatrics Microscopy Core for funding the Zeiss LSM800 confocal microscope.

## AUTHOR CONTRIBUTIONS

B.L.W. and R.K.M. conceived of the project. B.L.W. performed the experiments, analyzed data, and wrote the original draft of the manuscript. B.D.D. performed biochemical validation experiments and analysis for interaction data. Y.S. conducted structural modeling and analysis. B.L.W. and V.K.S. generated the windowed kidney live-imaging data. B.L.W., M.E.C., and A.R.M. performed experiments to generate knockdown and rescue data. R.K.M. oversaw the experiments and supervised the project.

## MATERIALS AND METHODS

### Xenopus laevis embryos

*Xenopus laevis* adult frogs were purchased from Nasco (LM00531MX and LM00713M) or Xenopus1 and maintained according to standard procedures. Eggs were manually collected from adult pigmented *Xenopus laevis* frogs approximately 14-20 hours post ovulation induction with human chorionic gonadotropin (hCG) injection. Collected eggs were fertilized *in vitro* using macerated sperm and placed in 0.3x MMR. Blastula orientation and stages were determined according to established methods (Brüser & Bogdan, 2017) and reared as previously described (Sive et al., 2000). All work was conducted in accordance with the University of Texas Health Science Center at Houston, Institutional Animal Care and Use Committee (IACUC) approved protocol #AWC-25-0074.

### Synthetic mRNA and Morpholinos

For mRNA injections, capped mRNA transcripts were synthesized from DNA-plasmids under standard protocols using an SP6 mMessage mMachine transcription kit (ThermoFisher, AM1340M) and purification. pCS2-membrane-tagged-RFP (mRFP) [Davidson et al., 2006], pCS2- membrane-tagged-EGFP (mEGFP) (Shindo & Wallingford, 2014), pCS2-Rab-11:GFP (Ossipova et al., 2014), and pCS2-β-galactosidase (Lyons et al., 2009; Miller et al., 2011) constructs were gifts from Dr. Raymond Keller’s lab, Dr. John Wallingford’s lab, Dr. Sergei Sokol’s lab, and Dr. Peirre McCrea’s lab respectively. pCS2-hDNMBP-GFP and pCS2-ΔN-hDNMBP-GFP constructs were generated using pcDNA3-HA-Tuba (Addgene plasmid 22214) to clone DNMBP (also known as Tuba) cDNA into BcuI/XbaI digested pCS2-GFP vector (cloning done by Epoch Life Science). Additionally, pCS2-hDNMBP-RFP, pCS2-hDNMBP-mKate2, pCS2-ΔN-hDNMBP-RFP, and pCS2- ΔN-hDNMBP-mKate2 constructs were also obtained from Epoch Life Science. Formerly developed translation-blocking morpholino Dnmbp MO1 (5’-TCGAACCACCGATCCCACCTCCATC-3’) (DeLay et al., 2019) and standard control MO (5′CCTCTTACCTCAGTTACAATTTATA 3′) were purchased from GeneTools, LLC (Philomath, OR, USA).

### Yeast two-hybrid

Screening using Human DAAM1 (aa 1-1077) as a N-LexA-DAAM1-C fusion, and then N-GAL4-DAAM1-C fusion employed a mouse kidney library in collaboration with Hybrigenics ULTImate Y2H. We obtained 2 positive clones over 88 million interactions tested for the LexA vector and 13 positive clones over 70 million interactions tested for the Gal4 vestor, which were then sequenced and analyzed.

### Embryo Microinjections

*Xenopus laevis* embryos were microinjected at the single-cell stage for Western blot analysis, or into the V2 blastomere at the eight-cell stage for embryonic kidney targeted microinjections (DeLay et al., 2016; Moody and Kline, 1990; Nieuwkoop and Faber, 1994). Injection solutions contained synthetic mRNAs alone, or in combination with antisense morpholino oligonucleotides (MOs). Single cell embryos were injected with 20 ng of MO for Western blot analysis and 8-cell embryos were injected with 10 ng of MO for phenotypic analysis. Concentrations of mRNAs co-injected with MOs, or injected alone, were as follows unless otherwise stated within context: memGFP [0.5ng], memRFP [0.5ng], hDNMBP-GFP, -RFP, or -mKate2 [250pg], ΔN-hDNMBP-GFP, -RFP, or mKate2 [0.5ng], E-cadherin-GFP [250pg], β-galactosidase [250pg].

### Immunoprecipitation

For *in vitro* pull-down assays involving N-Daam1 and C-Dnmbp, 5 µg of DNA encoding pCS2+HA-C-Dnmbp and pCS2+N-Daam1-Myc was separately transcribed and translated using the TNT SP6 High-Yield Wheat Germ Protein Expression System (Promega) per the manufacturer’s protocol. Synthesized proteins were then incubated together in three different tubes for IgG pull-down, Myc pull-down, and HA pull-down using amylose-resin.

For *in vivo* pull-down assays involving N-Daam1 and C-Dnmbp, one-cell stage embryos were co-injected with 500 pg of synthetic mRNAs for pCS2+HA-C-Dnmbp and pCS2+N-Daam1-Myc each. Lysates of stage 10-12 embryos for IgG, Myc, and HA were prepared as previously described (Cho et al., 2010). Protein-A/G agarose beads (Santa Cruz Biotechnology) were used to precipitate complexes prior to western blotting.

### Western Blots

Embryo lysates for various NF developmental stages (Nieuwkoop and Faber, 1994) were created from 10 pooled embryos of the same stage for each injection condition. Prior to collection, embryos were screened for presence of co-injected fluorescent injection tracer. Protein lysates were made as previously described (Kim et al., 2002). The equivalent of one embryo was used per lane of 4-20% precast polyacrylamide gel (Bio-Rad cat# 4561096). Proteins were transferred onto a PVDF or nitrocellulose membrane prior to KPL blocking.

### *Xenopus* Whole-Mount Immunofluorescence

*Xenopus laevis* embryos underwent kidney targeted microinjections at the 8-cell stage as previously described (DeLay et al., 2016) and were reared to tadpole stage 30-33 for early nephrogenesis or stage 38-40 for mature nephron analysis (Nieuwkoop and Faber, 1994). Embryos expressing fluorescent tracer were fixed with MEMFA (3.7% formaldehyde, 4mM MOPS, 2mM EGTA, and 1mM MgSO4, pH 7.4) for one hour at room temperature or overnight at 4°C. Following fixation, embryos were washed with 100% methanol (MeOH) twice over ten minutes and rehydrated through a series of phosphate buffered saline (PBS) containing 0.1% Triton X-100 (PBS-T) washes over one hour. Rehydrated embryos were incubated overnight in 20% goat serum diluted in PBS-T at 4°C to prevent non-specific targeting of primary antibody. For detection of Rab11, embryos were fixed with 2% trichloracetic acid solution (TCA) for 30 minutes at room temperature and washed in 0.3% Triton X-100 diluted in PBS over 30 minutes (Ossipova et al., 2014) prior to blocking as described above. Blocked embryos were incubated at 4°C overnight in 10% goat serum with primary antibodies. The following day, primary antibodies were recollected, and embryos were subjected to a series of 1X PBS-T washes for removal of excess antibodies. Primary antibody detection was achieved using fluorophore-tagged secondary antibodies against the host species for each primary antibody. Secondary antibody incubation in 10% goat serum took place at 4°C overnight. The following day, embryos were subjected to a series of 1X PBS-T washes to remove excess antibodies and dehydrated in 100% methanol, before being optically cleared with BABB/Murray’s clearing medium (1:2 volume of benzyl alcohol to benzyl benzonate) for confocal imaging. Embryos were identified for proper kidney-targeted injection prior to fixation and/or post staining using an Olympus SZX16 fluorescent stereomicroscope and kidney images were taken using a Zeiss LSM800 confocal microscope.

### Imaging of *Xenopus* Pronephros

*Xenopus* 8-cell stage embryos were microinjected with synthetic mRNA(s) into the V2 blastomere for kidney targeted injection, as described above. Embryos were reared to stage 28-32 (Nieuwkoop and Faber, 1994) and selected for positive kidney injection under a fluorescent stereomicroscope. To obtain high-resolution imaging of the nephric primordium *in vivo*, kidney-windowed embryos were created, as previously described by Krneta-Stankic et al., 2021. Removal of the surface epithelium to expose the developing nephron was accomplished using sharpened forceps (Fisher, NC9404145) under an Olympus SZX16 fluorescent stereomicroscope. Microsurgical procedures were performed in plastic petri dishes coated with 2% agar containing Danilchik’s for Amy (DFA) solution (53 mM NaCl, 5 mM Na2CO3, 4.5 mM Potassium Gluconate, 32 mM Sodium Gluconate, 1 mM CaCl2, 1 mM MgSO4, buffered to pH 8.3 with 1 M bicine) supplemented with 1 g/L of Antibiotic Antimycotic solution (1:100, Sigma, A5955). Live images were acquired using a Zeiss LSM800 microscope with Airyscan detector.

### Image processing and Statistical Analysis

E-cadherin localization between nephron progenitor cell junctions was detected by whole-mount fluorescence immunostaining and confocal microscopy. Junctional integrity was quantified by measuring the fluorescence intensity of E-cadherin along individual cell-cell borders using the Zen Lite image processing software and profiling tool. Analysis of E-cadherin intensity within the Dnmbp-KD kidney compared to the std-control kidney was conducted using Prism software.

### Structural Modeling

Structural modeling of the Daam1–Dnmbp complex was performed using AlphaFold3 (Abramson et al., 2024). The predicted structure was imported into UCSF Chimera version 1.19 (Pettersen et al., 2004) and subjected to energy minimization using the AMBER ff14SB force field (Maier et al., 2015) to remove local steric clashes while preserving the overall predicted conformation. Energy minimization consisted of 1,000 steps of steepest descent, followed by 1,000 steps of conjugate gradient minimization, both using a step size of 0.02 Å. The minimized model was used for structural visualization and analysis of the predicted interaction interface. Protein domains were annotated according to UniProt. Dnmbp domains included SH3-1 (aa 2–61), SH3-2 (aa 66–126), SH3-3 (aa 145–204), SH3-4 (aa 243–302), the DH domain (aa 784–967), the BAR domain (aa 1008–1217), SH3-5 (aa 1285–1348), and SH3-6 (aa 1513–1576). Daam1 contain three major domains, the FH3 domain (aa 45–420), FH1 domain (aa 528–599), and FH2 domain (aa 600–1009), with the actin-binding region located between amino acids 693 and 702 within the FH2 domain.

